# Stimulation of immunity-linked genes by membrane disruption is linked to Golgi function and the ARF-1 GTPase

**DOI:** 10.1101/2021.11.16.468795

**Authors:** Matthew J. Fanelli, Christofer M. Welsh, Dominique S. Lui, Lorissa J. Smulan, Amy K. Walker

## Abstract

Immunity-linked genes (ILGs) are activated by pathogens but also may respond to imbalances in lipids. Why pathogen attack and metabolic changes both impact ILG activation is unclear. We find that ILGs are activated when membrane phosphatidylcholine ratios change in secretory organelles in *C. elegans*. RNAi targeting of the ADP-ribosylation factor ARF-1, which disrupts the Golgi, also activates ILG expression, suggesting that activation of this membrane stress response could occur outside the ER. Our data argue that ILG upregulation is a coordinated response to changes in trafficking resulting from intrinsic cues (changes in membrane lipids) or extrinsic stimulation (increased secretion during immune response). Indeed, a focused RNAi screen of ILGs uncovered defects in secretion of two GFP reporters as well as accumulation of a pathogen-responsive CUB-domain fusion protein. These results also suggests that genes shared between the classical pathogen responses and lipid stress may act to counteract stress on secretory function.

**Teaser:** Pathogen response genes are also activated by lipid imbalances, which we suggest occurs because both processes put stress on the secretory pathway.

## Introduction

Cellular stress responses react to extrinsic insults by upregulating genes that defend the cell. Stress responses also restore organelle function in response to intrinsic changes. Several of these intrinsic stress response pathways are also linked to metabolic processes. For example, overnutrition can lead to metabolic changes that induce cellular stress through an increase in reactive oxygen species (*1*). Several signaling pathways play a dual role, functioning as coordinators of nutrient responses as well as stress responses. For example, the nutritional sensor mTORC1 can influence stress-responsive transcription (*2*). In addition, insulin signaling pathways are strongly linked to stress. In addition, mutations in *daf-2*, the *Caenorhabditis elegans* insulin receptor/insulin-like growth factor ortholog, produce highly stress-resistant animals (*3*). Finally, changes in ER lipids have a profound effect on stress pathways (*4–6*). Specialized ER sensors detect both the accumulation of unfolded proteins and overload of the secretory pathway. These sensors then initiate gene expression programs that reduce protein load or increase lipid production (*7, 8*).

While stress responses and metabolism are clearly intertwined, the biological advantages of linking these processes are less clear. For example, in a previous study, we found that RNAi-mediated knockdown of two different lipid synthesis modulators, *sams-1* and *sbp-1*, caused upregulation of pathogen-response genes in *C. elegans* (*9*). One of these lipid modulators, *sams-1/MAT1A,* encodes an S-adenosylmethionine synthase important for phosphatidylcholine (PC) production (*10*). The Troemel lab also identified *sams-1* as a regulator of the infection response gene *irg-1* in an RNAi screen (*11*). The other lipid modulator identified in our previous screen, *sbp-1/SREBPF1,* codes for a master transcriptional regulator of lipid synthesis genes (*12*). Reduction of *sbp-1/SREBPF1* causes a decrease in total lipid stores (*13*). Notably, mammalian *SREBPF1* knockdown also results in the enrichment of immunity-linked gene expression in human cells (*14*). Recent studies from other labs have found that ILGs are upregulated after mutation or RNAi of genes affecting PC synthesis. ILG activation was seen upon disruption of *pmt-2* (*15*) or *lpin-1* in the presence of excessive glucose (*16*). These immunity-linked genes were also upregulated in *skn-1* mutants, a transcriptional regulator of the stress response and metabolic genes (*17*). In addition, dietary restriction results in the upregulation of similar gene sets (*18*). These results together suggest that multiple types of lipid imbalance seem to impact immunity-linked gene expression and that these links may be conserved across species, as indicated by results in human cells cited above.

Exposure of *C. elegans* to bacterial pathogens stimulates the expression of a diverse set of genes, including antimicrobial peptides, enzymes for detoxifying xenobiotics, and neuromodulatory peptides to coordinate inter-organ defenses (*19–22*). However, many of the genes upregulated in response to either bacteria or fungi do not fall into these clearly defensive pathways (*23*) or have clear functional roles. Instead, these genes are defined by pathogen-responsive expression. In order to understand why knockdown of genes acting in lipid metabolism might activate genes linked to immune responses when no pathogens were present, we took two approaches. First, we looked for lipid signatures that could signal this response and were shared between *sams-1* and *sbp-1* by comparing lipidomics of whole animals and fractioned extracts. Second, we performed a RNAi screen of genes functioning in complex lipid synthesis and ER/Golgi dynamics for activation of the immunity-linked reporter, p*sysm-1::GFP* (*24*). In the lipidomics studies, we found that *sams-1 and sbp-1* knockdown animals had broad and distinct changes in their lipidome, but both showed lowering of PC levels in the ER/Golgi fraction which we previously linked to SBP-1/SREBP1 maturation in *sams-1* animals (*25*). Strengthening the links to PC, our targeted RNAi screen showed that synthesis of PC, increased p*sysm-1::GFP* expression. RNAi of Golgi/ER transport regulators, including the GTPase *arf- 1/ARF1,* activated the immunity-linked reporter. Notably, some ILGs were also upregulated after *arf-1*/ARF1 RNAi, suggesting that disrupting Golgi function regulates immunity-linked genes. Finally, we performed a targeted screen, and found that interfering with ILGs expression in *sbp-1(ep79)* animals disrupted trafficking of a secreted reporter. Taken together, our results suggest that some ILGs act to balance the increased secretory load in pathogen-stimulated cells or when membrane lipid balance is perturbed. While immune function is critical for host defenses, immune activation in non-pathogenic conditions, such as in metabolic disease, may have deleterious effects (*26*). Therefore, it is critical to define cellular roles for pathogen-responsive genes. Our studies suggest that genes upregulated in response to both pathogens and lipid perturbation in *C. elegans* could act to support secretory processes altered in both biological contexts.

## Results

### Gene set enrichment shows upregulation of pathogen-response genes in multiple models of membrane dysfunction

Levels of lipids within membranes are tightly monitored, and imbalances may induce cellular stress pathways that regulate genes to restore lipid ratios (*8*). We previously found that *sams-1*, *lpin-1,* and *sbp-1* are part of a feedback loop that responds to shifts in membrane PC levels and activates immunity-linked genes (**Fig. 1A**)(*9, 25*). In order to test why immunity-linked genes are activated after disruption of lipid-synthesis regulators we first conducted a bioinformatics analysis using our recently developed *C. elegans* specific gene ontology tool, WormCat 2.0. This computational analysis allows combinatorial graphing of gene enrichment scores based on detailed annotation of all *C. elegans* genes (*27, 28*) and uses a nested annotation strategy (Category1/Category2/Category3) to provide broader or more detailed information on genome-wide datasets. The WormCat annotation strategy also includes a specialized category, Unassigned, to include genes cannot be classified using our strict criteria. Genes that are generally characterized are placed in UNASSIGNED at the Category 1 level in WormCat 2.0 (“other” in the 1.0 version), but we also use unassigned to denote genes within larger, well described category whose specific roles were less clear. For example, in WormCat, genes with roles in immunity are classified at the Cat1 and Cat2 level in STRESS RESPONSE: Pathogen. This category is broken down into Cat3 classifications for transcriptional or signaling regulators, neuropeptide-like proteins, proteins containing common antimicrobial motifs (saposin, caenacin, CUB, and others).

**Figure 1:**
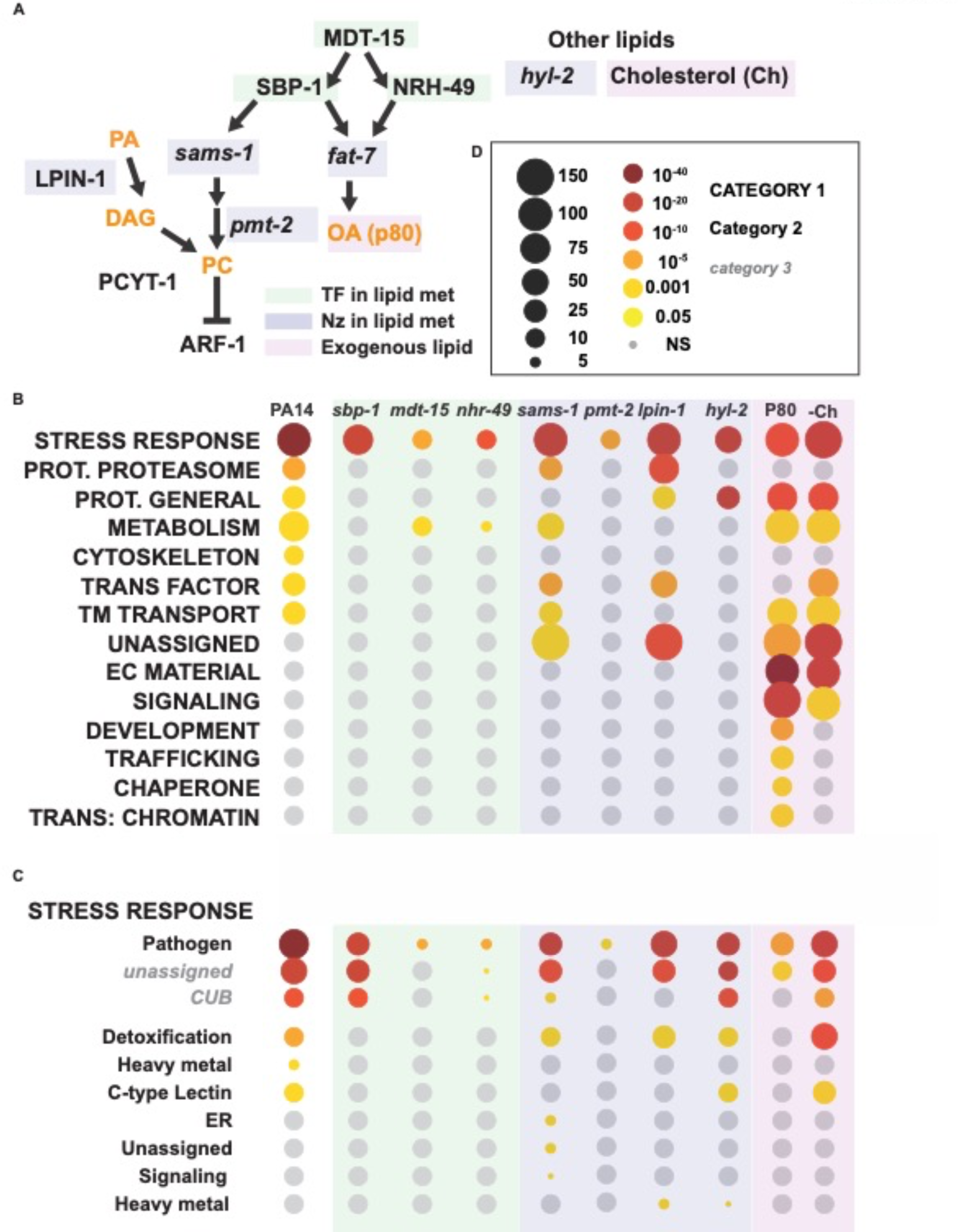
Upregulation of immunity-linked genes occurs downstream of multiple types of membrane lipid disruption. **A.** Schematic diagram of lipid metabolic genes linked to immune gene expression. Meta-analysis of published RNA seq data from linking lipid perturbation to immune gene expression using WormCat 2.0 for Category 1 (Cat1) (**B**) and for the Category 2 (Cat2) level under STRESS RESPONSE (**C**). Legend is in **D.**

This category also includes 97 genes upregulated during the pathogen response, but lacking molecular characterization, which were designated STRESS RESPONSE: Pathogen: *unassigned* (*27, 28*). This sub-category had the highest enrichment after RNAi of *sbp-1* or *sams-1* (*27–29*), suggesting that immunity linked genes upregulated after lipid unbalance could function in area common to immune response and lipid stress.

In order to better characterize these ILGs, we used WormCat to perform a meta-analysis of published data comparing categories enriched in genes upregulated by the pathogen PA14 (*22*) with multiple models of disrupted lipogenesis (**Fig1A**). We found enrichment in Category1(Cat1) STRESS RESPONSE, and more specifically in Pathogen at the Cat2 level in genes upregulated after the lipogenic transcriptional regulators *sbp-1* (*9*), *mdt-15* (*30*) or *nhr-49* or enzymes acting in lipid synthesis, *sams-1* (*9*), *lpin-1* ((*16*); **DataS1**), *pmt-2* (*6*), and *hyl-2* were knocked down as well as when exogenous lipids such as P80 (oleic acid mimic) (*33*) or cholesterol were added (*34*) (**Fig1B-C)**. Notably, the strongest enrichments at the Cat3 level were in STRESS RESPONSE: Pathogen: *unassigned*. Genes in this category are defined by solely upregulation upon pathogen exposure in *C. elegans* but do not have defined functions as antimicrobial peptides or within pathogen-stimulated signaling pathways such as MAPK signaling (*27, 35*). Importantly, this class of genes is not explicitly defined by a traditional GO analysis. Genes upregulated in *pmt-2(RNAi)* animals were also linked to the Lipid Bilayer Stress induced Unfolded Protein Response (UPR^LBS^), a UPR^ER^ mechanism (*6*), therefore, we compared WormCat enrichment of genes increasing after tunicamycin treatment (*36*). We found that while STRESS RESPONSE: Pathogen: *unassigned* genes were enriched, the category enrichment was independent of the ER stress regulators *xbp-1* or *ire-1 (***FigS1A)**, suggesting potential for a distinct regulatory mechanism.

We also noted that the landscape of upregulated genes was complex across other categories, with STRESS RESPONSE: Detoxification enrichment scores increasing in both *sams-1* and *lpin-1* but not *sbp-1* (**Fig1C**). In addition, METABOLISM: Lipid was increased after *lpin-1* RNAi (**Fig1C**). These data are consistent with our previous studies showing that SBP-1 transcriptional targets are upregulated in these animals (*10*). Interestingly, immunity-linked categories after SREBP-1 knockdown in human melanoma cell lines (*14*). Thus, genes linked to immunity are upregulated across a diverse set of lipid synthesis modulators in *C. elegans* and mammalian cells and occur in the absence of pathogen stimulation.

SBP-1/SREBP1 is a basic helix-loop-helix transcription factor required to express a suite of lipid metabolic genes, including *fat-7* in *C. elegans* and its ortholog SCD1 in mammals (*12, 13*). It is also necessary for the upregulation of *fat-7* in lipid-replete *sams-1(lof)* animals ((*10*) (**FigS1B**). Since SBP-1 upregulates lipid synthesis genes, we tested the possibility that it directly functions in ILG upregulation by using *sams-1(RNAi)* to increase the expression of multiple SBP-1 target genes (*10*) and assessing the need for *sbp-1*. As we previously reported in Walker, 2011, *fat-7* depends on *sbp-1* in both the control and *sams-1(RNAi)* background. However, the ILG *hpo-6* is expressed at high levels when induced by reduction in *sbp-1*, *sams-1*, or in *sams-1(lof); sbp- 1(RNAi)* animals (**FigS1C**), suggesting that the effects of SBP-1 on ILG upregulation are indirectly related to its function as a transcriptional regulator.

Many ILGs in *C. elegans* act downstream of signals from the p38/MAPK14 kinase PMK-1 (*22*). In previous studies in *C .elegans*, we found that low PC in *sams-1(lof)* animals leads to constitutive phosphorylation of PMK-1 and that ILG induction was dependent on both PC and *pmk-1* (*9*). We confirmed that knockdown of *sbp-1* also induces PMK-1 phosphorylation (**FigS1D**). Since ILGs can be induced in mammalian cells in response to reduction in *SREBPF1*, we used siRNA to knock down the rate-limiting enzyme for PC production, *PCYT1,* in human hepatoma cells (**FigS1E**). We found that p38/MAPK14 phosphorylation is induced to levels similar to lipopolysaccharide (LPS). Taken together, the results suggest that upregulation of ILGs in response to lipid perturbations may share regulatory link to pathogen-stimulated pathways.

### Loss of sbp-1 decreases PC levels in ER/Golgi extracts

Immunity-linked gene expression and PMK-1 phosphorylation both occur after reduction in *sams-1* and *sbp-1*, although these genes act at different points in lipid synthesis regulation. In order to identify shared lipid signatures that might stimulate this process, we analyzed the total lipidomes of these mutants by LC/MS. Because animals were lysed directly into lipid extraction buffer, levels were normalized to total lipid as in (*25*). The entire lipidome was broadly altered in both cases, with 27% of lipid species showing significant changes in *sams-1(RNAi)* animals and 30% after *sbp-1(RNAi)* (**Fig2A, B; DataS2**). We next examined lipid levels at the class and species level. As in our previous studies (*10*), *sams-1* animals showed lower PC and higher TAG (Triacylglyceride) (**Fig2C**), and we also noted lower phosphatidylserine (PS) and sphingomyelin (SM) levels in comparison to other lipid classes (**Fig2C, FigS2C**). TAG was lower after *sbp-1* RNAi, as expected from previous studies (*37*). We also found that while reductions in PC did not reach significance, PE (Phosphatidylethanolamine) increased, and Cer (Ceramide) levels dropped (**Fig2C, D**). In short, *sbp-1* RNAi induces broad changes in the lipidome across multiple classes of lipids.

**Figure 2:**
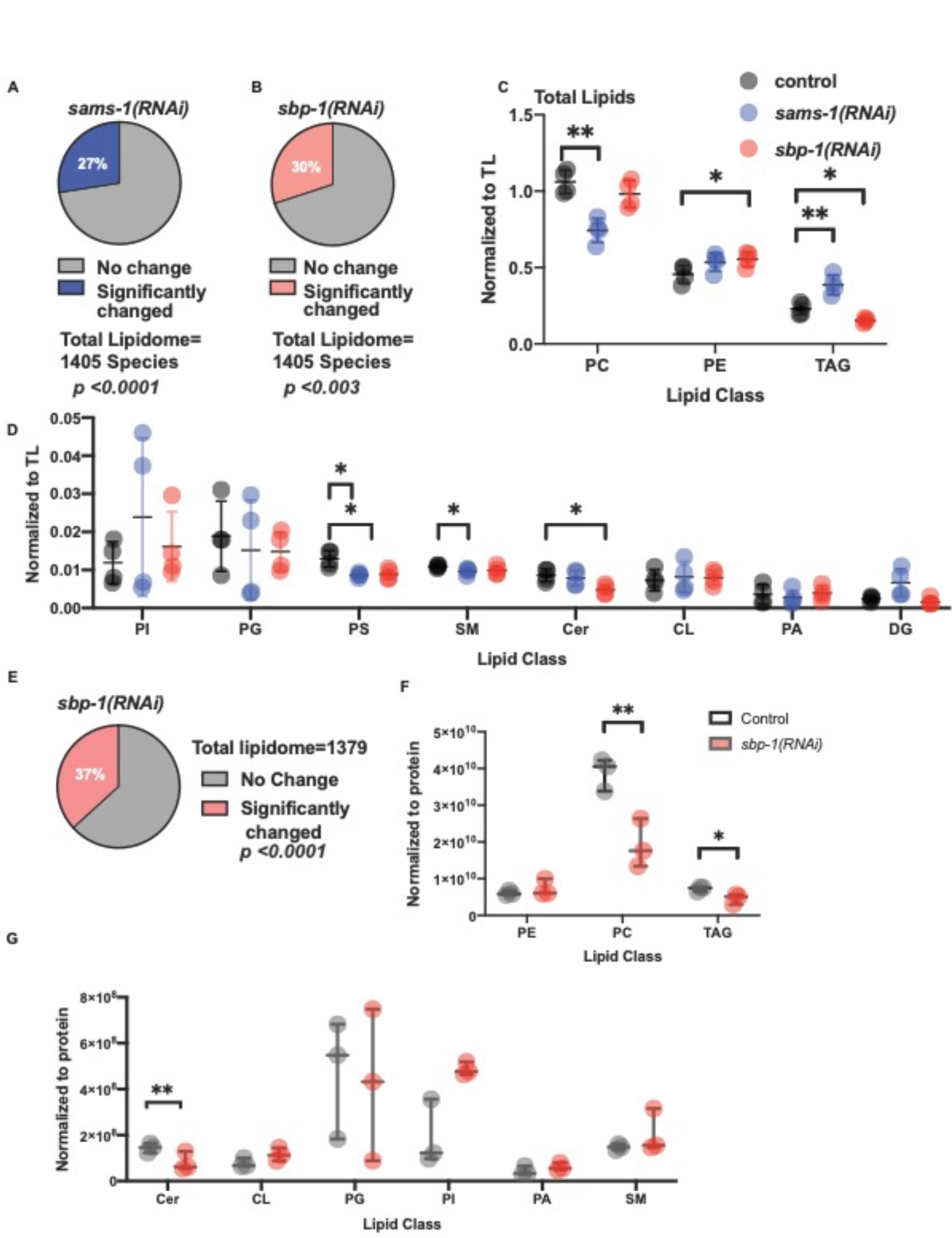
Broad lipidomic changes in total and microsomal lipid compartments after *sams- 1* or *sbp-1* RNAi. LC/MS lipidomics show that nearly a third of total lipid species change after *sams-1* or *sbp-1* RNAi (**A, B**). Significant changes in major lipid classes are mostly distinct in *sams-1* and *sbp-1* RNAi (**C, D**). See also **Figure S1** (**B, D**) for lipid species distributions and additional lipid class analysis and **Data S2** for all values. The microsomal (ER/Golgi) lipidome is also broadly altered after *sbp-1 RNAi* (**E**) and shows changes in different lipid classes than in the total lipidome (**F, G**). Error bars show standard deviation. Lipid class data is calculated by students T-Test. See Figure S1 for lipid species distributions and additional lipid class analysis.

The properties of lipid classes can be altered when distributions of acyl chains alter membrane properties (*38*). We find that there are significant differences in the populations of species within phospholipids (PC, PE, lyso-phosphatidylethanolamine (LPE), and phosphatidylglycerol (PG), TAG, diacylglycerides (DG), and SMs after *sams-1(RNAi)*. Changes in species distribution in lipid classes after *sbp-1* RNAi were limited to TAG, lyso-phosphatidylcholine (LPC), and Cer in agreement with previous studies showing lower TAGs in *sbp-1(RNAi)* animals. We asked if *sams-1* and *sbp-1* RNAi caused similar shifts in animals and found overlapping changes in 180 shared lipid species (**DataS2**). Fifty-seven of these species were decreased in both, 17 increased in both, and 102 lipid species changed in opposite directions out of 1405 total lipid species detected. Lipid species diverged most strongly within the TG class, with increases in *sams- 1(RNAi)* animals and decreases after *sbp-1* RNAi (**DataS2**). This agreed with total class levels. For lipid species that increased in both animals, PE species were the highest, with 15 changing at significant levels. Seventy-four PC species changed in both *sams-1* and *sbp-1* knockdown animals, with 35 decreasing in both (**TableS3**). Finally, three SM species decreased in both animals. Thus, while *sams-1* or *sbp-1* knockdown each lead to broad changes in the total lipidome, relatively few overlapping species changed in both instances. Of those, PC species were the best represented among lipid species that decreased upon knockdown of either *sams-1* or *sbp-1* in animals.

The total lipidome contains multiple organelles, including the plasma membrane, mitochondria, nuclear membrane, and Golgi/ER. However, lipid ratios in specific organelles may be reflective of particular functions; for example, we previously found that PC ratios in ER/Golgi fractions were associated with activation of SBP-1 in *sams-1* and *lpin-1* animals (*25*). Therefore, we performed LC/MS lipidomics on microsomes from *sbp-1(RNAi)* animals to assess lipids in the ER/Golgi fraction. Because the lipidomics was performed on fractionated animals, we used total protein for normalization. We found a broader fraction of the ER/Golgi lipidome was affected after *sbp-1(RNAi)* than in the total lipid samples. Nearly 40% of lipid species were altered in these fractions (**Fig2E; Data23**), with significant changes in species across most classes (**FigS2C**). Similar to the unfractionated extracts, levels of TAG and Cer decreased in the ER/Golgi (**Fig2F**). Notably, significant ER/Golgi-specific decreases occurred in PC and DG (**Fig2F; FigS2D**). This demonstrates that *sbp-1(RNAi)* has distinct effects on the ER/Golgi membranes and shows decreases in PC levels at the sub-compartment level, similar to *sams-1* RNAi.

### Targeted RNAi Screen Reveals roles for COP I transport in immune gene expression

Low PC levels are associated with ILG expression in *sams-1* (*9*) and *pmt-2(RNAi)* animals (*6*). The low PC in the ER/Golgi compartment of both *sams-1* (*25*) and *sbp-1* RNAi animals suggests this could be a shared signature linked to ILG upregulation. To understand how these lipids might act, we next sought to identify genes important for ILG expression. We screened for upregulation of an immune reporter, p*sysm-1*::GFP (*24*), with an RNAi sub-library that targets genes involved in complex lipid metabolism, lipid signaling and selected ER/Golgi transport regulators. We previously used a lipid-function RNAi sub-library to identify low-PC regulators of SBP-1/SREBP-1 (*25*). Notably, we constructed RNAi clones for genes not represented in the Arhinger or Orfeome RNAi libraries. The resulting library includes all predicted phospholipases and enzymes for synthesis of complex lipids (*25*). For the current screen, we expanded the sub-library to include more genes involved in ceramide/sphingomyelin synthesis and confirmed all clone identities by sequencing. Thus, this library has a broader representation of genes linked to complex lipid synthesis than the commercially available RNAi libraries. p*sysm-1*::GFP is strongly induced by *sbp-1(RNAi)* (**Fig3A**) and has been used by multiple labs as a robust marker for immune gene induction (*18, 39, 40*).

**Figure 3:**
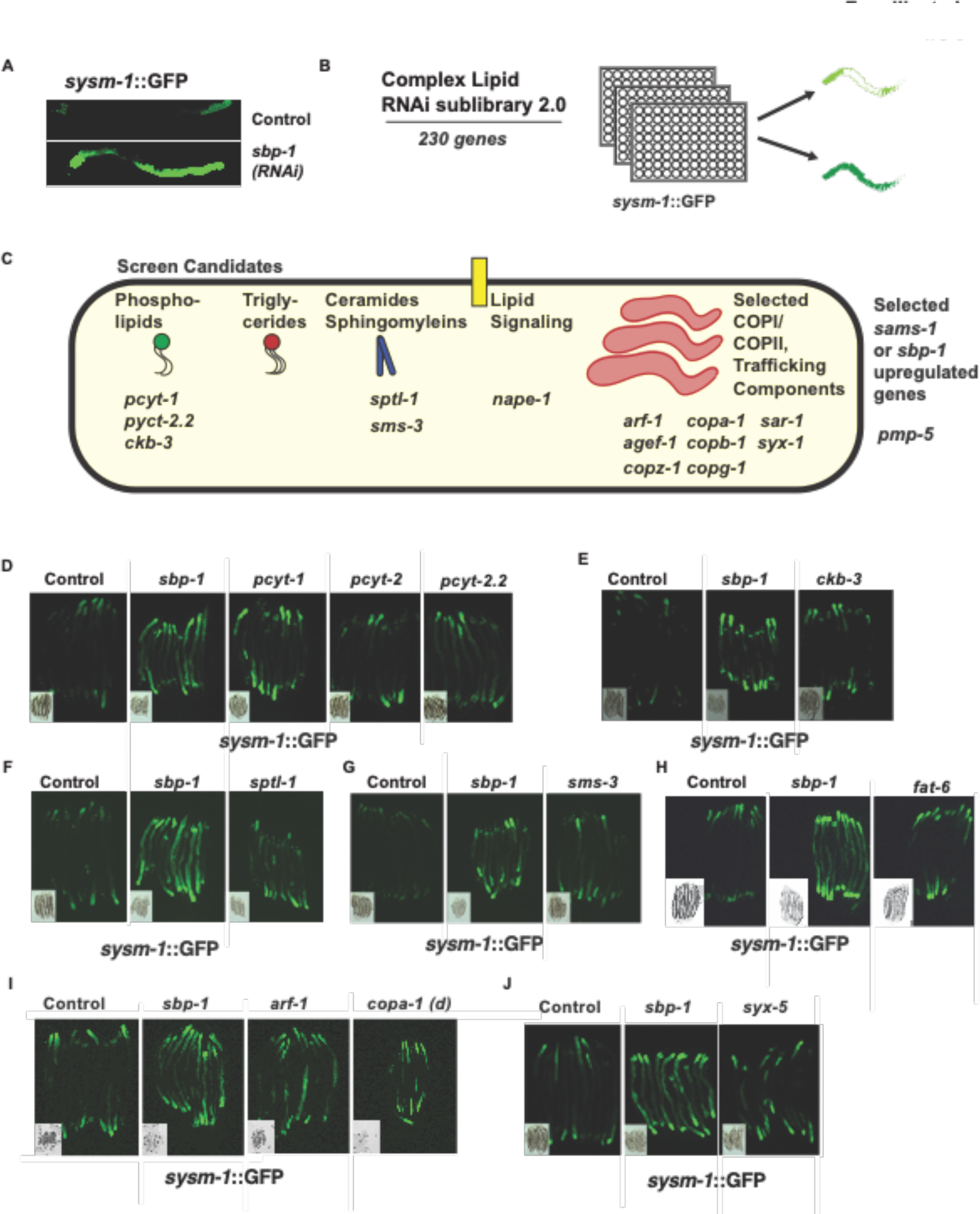
Targeted RNAi screen reveals candidates linked to Golgi/ER trafficking and ceramide/sphingomyelin synthesis impact immunity gene expression. (**A**) Immune gene reporter p*sysm-1*::GFP is increased after *sbp-1* RNAi. (**B**) Screen schematic. (**C**) Schematic showing RNAi library coverage and screen candidates. Epifluorescence imaging showing validation of reporter activity after RNAi of lipid PC synthesis genes (**D, E**), SM synthesis genes (**F, G**) and the stearoyl CoA desaturase *fat6/7* (**H**). Epifluorescence images showing GFP levels after RNAi of the *arf-1* GTPase or coatomer component *copa-1* or proteins acting at the Golgi such as *syx-1*/Syntaxin5 (**I, J**). Because of poor larval development, bacteria for *copa-1* RNAi was diluted to 1:10. Internal control for effect of *sbp-1(RNAi)* included in each panel and DIC image included as an inset for each RNAi. Quantitation of images is in **Figure S4, A, B**.

In order to identify genes in lipid synthesis important for ILG upregulation, we screened this library in quadruplicate in p*sysm-1*::GFP animals and identified a list of 20 candidates for retesting (**Fig3B; DataS3**). Retesting to identify the strongest candidates occurred in 3 stages: visual rescreening (4X), imaging of candidates from the rescreen, and qRT-PCR to assess changes in GFP expression. We also included the *sbp-1* target genes *fat-5*, *fat-6,* and *fat-7* (*13, 37*) in the visual retesting. These genes are orthologs of mammalian Stearoyl CoA desaturases (SCDs) (*41*) and can change membrane fluidity (*42, 43*) or induce lipid bilayer stress by changing ratios of saturated/unsaturated acyl chains within lipid classes (*6, 15, 44*).

Candidates from the targeted screen fell into two major groups. First, enzymes linked to PC and SM synthesis were found (**Fig3C-G; FigS3A, B; DataS3**). This included two isoforms of the rate-limiting enzyme for PC production, *pcyt-1,* and *sptl-1*, which initiates the first committed step in sphingolipid synthesis. Thus, our genetic data supports the notion that changes in PC or SMs are a shared signature linked to ILG activation in *sams-1* and *sbp-1* RNAi animals.

Notably, *fat-5, fat-6,* or *fat-7* RNAi, which would change acyl chain saturation within each lipid class, had modest effects on the p*sysm-1*::GFP immune reporter (**Fig3H, FigS3A, B; DataS3**), suggesting changes in membrane fluidity downstream of these enzymes is not a major contributor to the immunity-linked gene upregulation.

Second, Golgi/ER trafficking genes included four genes forming the COPI complex (which also caused developmental delays) and ARF1 guanine activating factor (*agef-1*). The ARF-1 GTPase, a key regulator of these factors, was a false negative in the original screen, but strongly activated the reporter in the rescreening. (**Fig3I**; **DataS3**). The coatomer proteins work in a complex (*45*) and are required for viability (*46*). Therefore, we chose one candidate, *copa-1,* and performed RNAi with diluted bacteria, allowing animals to develop fully. We found that the immune activation reporter was strongly expressed in the *copa-1* knockdown (**Fig3I**).

Stimulation of the reporter after the loss of multiple parts of the COP I machinery and its key regulator *arf-1* strongly argues that ER/Golgi dynamics signal to induce immunity-linked genes. The p*sysm-1*::GFP reporter represents a single gene in the immune response program. To more broadly survey ILGs in our screen validation, we used qRT-PCR with primers specific to *irg-1, irg-2,* and *hpo-6*, and found robust activation of each of our candidate reporters after knockdown of *arf-1* or *copa-1*, but more modest effects with the SM genes or SCDs (**FigS4C-H),** implicating ARF1 and Golgi function in ILG activation.

### Disruption of ER/Golgi activates immunity-linked genes

Cycling of the ARF1 GTPase requires interaction with the Golgi membrane (*45*). Alteration of membrane lipid ratios can affect interactions with ARF1 regulatory proteins, limiting GTPase cycling and blocking ARF1 activity (*25, 47*). We previously linked changes in PC levels in *sams-1* microsomes to the inactivation of ARF-1, which drove proteolytic processing of SBP- 1/SREBP-1 (*25*). Therefore, we reasoned that multiple modes of membrane lipid disruption could signal secretory stress through ARF-1 and activate ILGs. The upregulation of *sysm-1* and other immune genes after *arf-1* RNAi prompted us to examine genome-wide changes in the transcriptome that could occur in response to the loss of ARF-1 function. Using RNA-seq, we found that 705 genes were significantly upregulated after *arf-1* RNAi (**Fig4A, DataS4**) and that 40% of genes were also upregulated in *sams-1* or *sbp-1* RNAi animals (**Fig4B**). Downregulated genes in *sams-1, sbp-1,* or *arf-1* animals had low levels of overlap (**Fig4C**). WormCat gene set enrichment shows that STRESS RESPONSE: Pathogen: *Other* gene sets are significantly upregulated in *arf-1(RNAi)* animals as they are after *sbp-1, sams-1,* or *lpin-1* RNAi (**Fig4D-F**). Next, we examined WormCat enrichment within the genes that were shared between these three RNAi sets and found that shared genes retained strong enrichment for STRESS RESPONSE: Pathogen (**Fig4D-F**). Thus, *arf-1* GTPase knockdown causes similar or overlapping effects in immune gene expression to those seen after broad lipid disruption by *sbp-1* RNAi.

**Figure 4:**
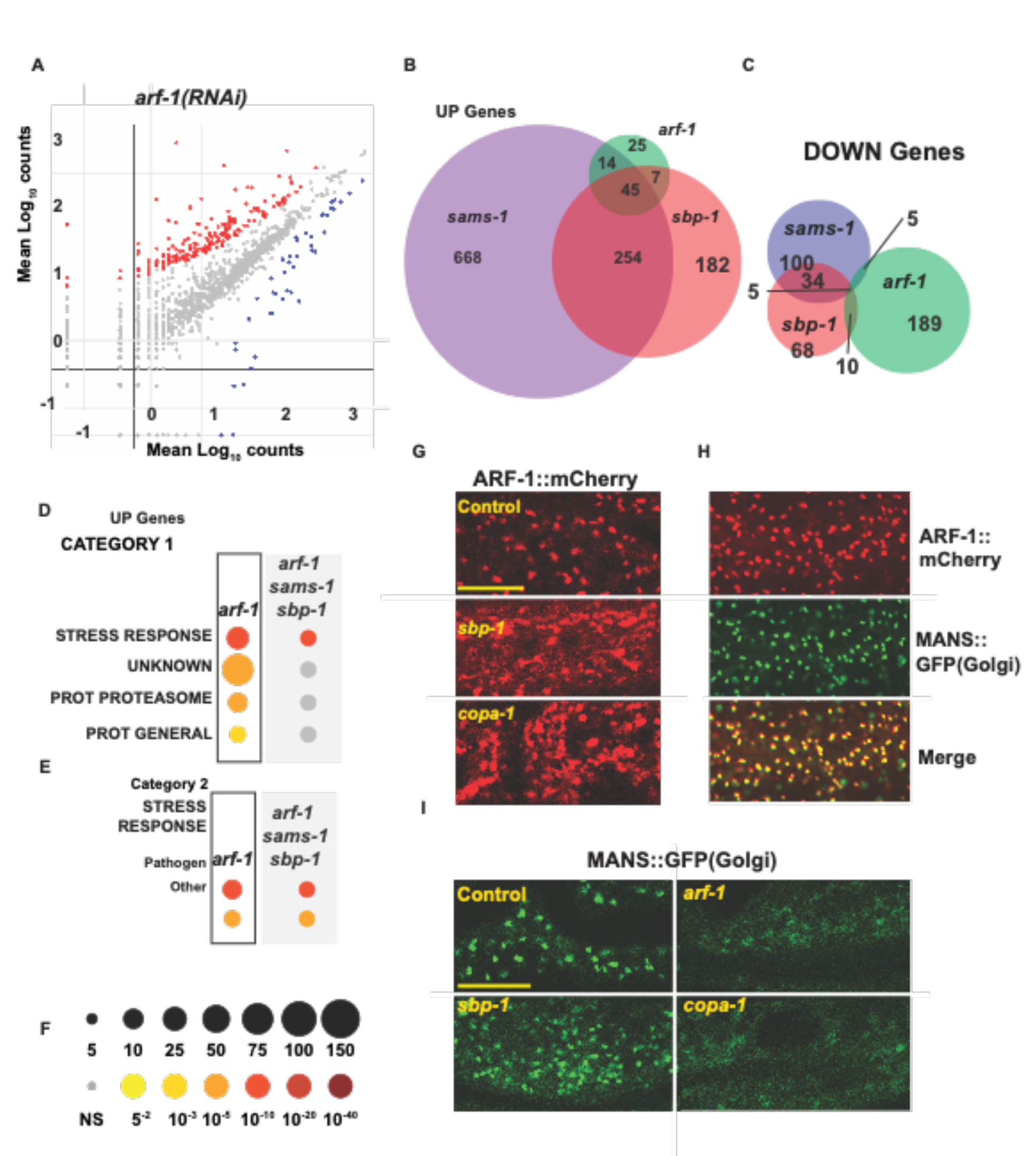
Disruption of Golgi/ER trafficking increases expression of immunity-related genes. **A.** Scatter plot shows changes in gene expression after *arf-1* RNAi. Venn diagrams showing limited overlap of up- (**B**) or downregulated (**C**) genes after *sbp-1, sams-1* or *arf-1* RNAi. WormCat gene set enrichment chart shows an increase in stress response and pathogen linked genes in Category1 (**D**) and Category2 levels (**E**). Legend is in **F**. **G.** Confocal projections of endogenously tagged ARF-1::mCherry reveal that that Golgi-like pattern is disrupted after *sbp-1* RNAi and quantitation in **Figure S5A**. **H**. Confocal images of intestinal cells from ARF-1::mCherry; ges-1::MANS-1::GFP animals. Scale bar is 10 microns. **I**. Confocal projections of the *C. elegans* Golgi marker MANS::GFP (*49*) in Control, *sbp-1, arf-1,* and *copa-1 (1:10)* RNAi intestines with quantitation of puncta intensity and number in **Figure S5B, C**. Scale bar is 10 microns. Significance is determined by the Mann-Whitney test.

In order to study the localization of ARF-1, we obtained a strain where endogenous ARF-1 is C- terminally tagged with mCherry by CRISPR (WAL10). Next, we asked if ARF-1 localization was altered in *sbp-1(RNAi)* animals by examining expression in the ARF-1::mCherry animals. In intestinal cells, ARF-1::mCherry formed a punctate pattern (**Fig4G-H**) characteristic of the Golgi ministacks occurring in *C. elegans* cells (*48*) and which localize adjacent to a Golgi marker, MANS::GFP (*49*), expressed in the intestine. MANS::GFP is a fusion of *aman-2* (alpha-mannosidase, a conserved Golgi specific protein) and marks the mini-Golgi stacks characteristic of these of *C. elegans* (*50*). This suggests that, as in other eukaryotes, *C. elegans* ARF-1 acts at the Golgi apparatus. Strikingly, ARF-1::mCherry appeared to form larger, more irregular punctae in *sbp-1(RNAi)* animals as well as having additional diffuse localization, similar to patterns seen with RNAi of the coatomer protein COPA-1 (**Fig4C**). This suggests that ARF-1 functions at the Golgi are altered in *sbp-1(RNAi)* animals.

Loss of human ARF1 or blocking ARF1 cycling with the fungal toxin Brefeldin A disrupts Golgi integrity (*51*). In addition, our previous studies found that lowering PC levels through RNAi of *sams-1* or knockdown of *PCYT1* (the rate-limiting enzyme for PC production) in mammalian cells blocked ARF1 GTPase activity and disrupted Golgi structure (*10, 25*). Because PC levels in *sbp-1(RNAi)* ER/Golgi fractions decreased (**Fig2C**) and ARF-1::mCherry was mis-localized, we next examined Golgi structure after *sbp-1(RNAi)* in animals where MANS::GFP was driven by an intestinal reporter (*49*). Consistent with previous studies (*52, 53*), knockdown of *arf-1* dramatically shifts the Golgi puncta to diffuse localization across the cytoplasm (**Fig4I; Fig S5A**). *copa-1* RNAi also diminished Golgi stacks along with altering ARF-1::mCherry localization. *sbp-1* knockdown affects Golgi structure, but results in a different pattern, in which Golgi stacks are smaller and more numerous with increases in diffuse cytoplasmic localization (**Fig4I, Fig5A**). We also examined MANS::GFP patterns in other candidates from our screen and noted that like *arf-1*, *copa-1* and *sar-1* were required for MANS-1::GFP puncta (**FigS5B, C**), as expected for core constituents of COP I and COP II transport. Contributors to PC synthesis also broadly abrogated MANS::GFP puncta, however, change in fatty acid desaturation, which induces UPR^LBS^ (*6*), caused a slight increase in MANS::GFP puncta intensity, suggesting that membrane stress initiated by Golgi misfunction has separable defects. Our screen candidates also included genes from ceramide synthesis, which can also affect membrane function (*54*). Notably, *sptl-1(RNAi)* did not noticeably alter Golgi marked by MANS::GFP and ARF- 1::mCherry puncta were visible (**FigS5A, D**), suggesting that *sptl-1* knockdown may mediate effects on immune genes through distinct cellular membranes. Taken together, we find that ARF-1 activity (*25*) and localization (this study) can be affected by *sams-1* or *sbp-1* RNAi and that targeting *arf-1* or the coatomer components can activate ILGs. Thus, blocking or limiting ARF-1 function at the Golgi may be part of the signal to activate these genes.

**Figure 5:**
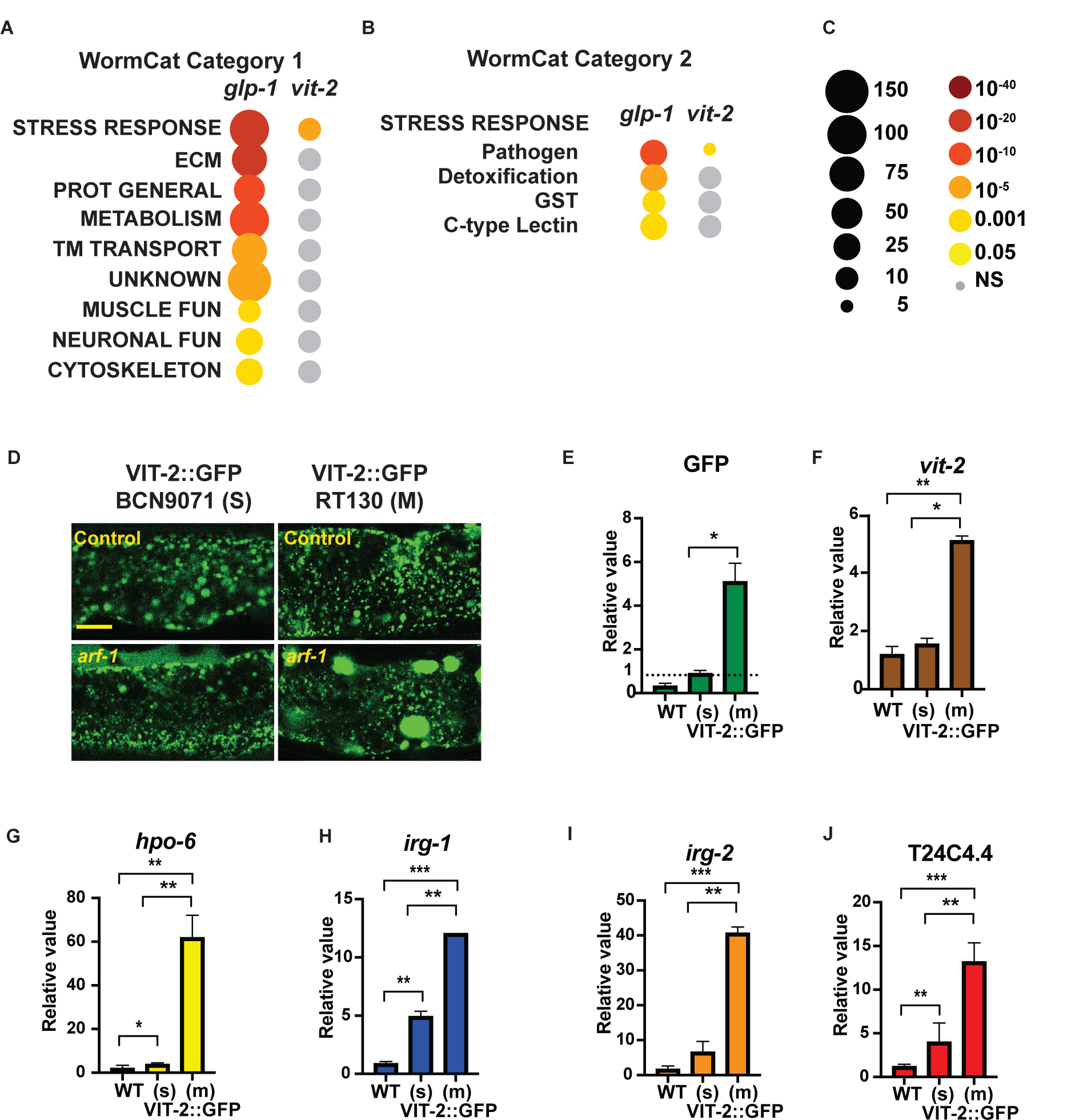
Immunity-linked gene expression is linked to trafficking disruption. WormCat gene enrichment of upregulated genes from *glp-1*(AA2735) (*63*) and *vit-2(ac3)* compared to SJ4005 (*64*) for Category 1 (**A**) and Category 2 (**B**). Legend is in **C**. **D.** Confocal projections of *C. elegans* intestinal cells expressing *vit-2*::GFP (RT130) or *vit-2*::GFP (*BCN9701*) in wild type or *arf-1(RNAi)* animals. qRT-PCR comparing gene expression in wild type, single copy *vit- 2::GFP* (BCN9071, s) or multi copy (RT130, m) animals. **E** and **F** show increase in GFP or *vit-2*. Immune response genes are shown in **F-J**. Significance is determined by Students T Test.

### Immunity-linked gene upregulation in models of trafficking dysfunction

The Golgi apparatus accepts proteins from the ER that are destined for secretion, processing them by glycosylation before secretory vesicles are loaded (*55*). Stimulation of the innate immune system may significantly impact trafficking load, as antimicrobial peptides are shuttled through the membrane trafficking system (*56*). Indeed, activation of the ER stress response has been noted in multiple systems when large numbers of proteins need to be produced and secreted (*57*). Interestingly, previous data suggest that induction of membrane stress may also be a sensor of infection (*58*). Based on work by the Ewbank lab, they also suggested that some pathogen-response genes act to support the trafficking of antimicrobial peptides (AMPs) (*58, 59*). We noted that the immunity-linked genes upregulated at low PC are largely outside of the antimicrobial categories and are comprised of gene sets defined by their shared expression upon pathogen exposure rather than function (*23, 27*). We hypothesized that some of these genes might be responding to direct stress on the trafficking system.

To explore this idea, we turned to two commonly used systems for studying trafficking in *C. elegans*, comparing ILG gene expression in strains with increasing copy number of VIT-2::GFP (*60*) and visualization of trafficking in both VIT-2::GFP and *myo-3*::ssGFP (*61*). *vit-2* is a vitellogenin produced and secreted from intestinal cells then taken up by the germline (*62*). *myo-3*::ssGFP has a signal sequence on the fluorophore and is expressed in body wall muscle (*61*). First, we used WormCat to examine category enrichment data from two published studies examining genome-wide mRNA expression patterns when *vit-2* is overexpressed or misregulated, placing stress on the secretory system. The Blackwell lab showed that germline- less *glp-1* mutants exhibit *vit-2*::GFP buildup near its production site in the intestines (*63*). WormCat analysis of their RNA seq data found strong enrichment patterns in STRESS RESPONSE: Pathogen. (**Fig5A**, **C; DataS4**). Next, we used WormCat to examine RNA-seq data from VIT-2::GFP (*64*) overexpressing animals produced by the Aballay group and also found increases in the Stress Response category enrichment (**Fig5B, C; DataS4).** As two models sharing VIT-2::GFP secretory overload show increases in STRESS RESPONSE: Pathogen genes, we sought to directly test this model by comparing ILG expression in two strains with a differing copy number of *vit-2*; RT130, made by microparticle bombardment and containing the GFP array in addition to wild type copies (*65*) and BCN9071 which is a CRISPR- generated allele integrated into the endogenous locus (*66*) (**Fig5D-F**). While VIT-2::GFP intestinal puncta appeared similar in control conditions, RNAi of *arf-1* produced larger aggregates in RT130 (**Fig5D**), suggesting that this strain is sensitized to stress in the secretory system. Strikingly, expression of ILGs increase with VIT-2::GFP copy number (**Fig5G-J**), demonstrating that ILG expression can be induced in the absence of pathogens by altering increasing expression of proteins destined for the secretory pathway.

### Knockdown of STRESS RESPONSE: Pathogen genes disrupt a trafficking reporter

We found that ILGs become upregulated in the absence of pathogen response when lipid levels become unbalanced, ARF-1 functions are compromised, or when production of secreted proteins increases. This prompted us to ask if some ILGs might function as part of a response to help stimulate passage through the secretory pathway when trafficking load is high, as when pathogen responses drive antimicrobial protein production, or when the structures of the secretory system are compromised. As a pilot experiment, we focused on one of the most strongly upregulated genes, *hpo-6*, which is not present in any of the commercially available RNAi libraries. To study its function more closely in this context, we made a cDNA construct to allow RNAi in VIT- 2::GFP RT130 in wild type and *sams-1(lof)* animals (*9*). *hpo-6* was originally identified in a screen for genes whose loss increased sensitivity to a pore-forming toxin (*67*). It has a glycoprotein domain and may occur in membrane rafts (*68*), but has no apparent homology to human proteins. *hpo-6(RNAi)* causes a slight increase in VIT-2::GFP in wild-type animals (**FigS6A**). However, in the low PC *sams-1(lof)* background, VIT-2::GFP pooling and aggregation suggested a broad effect on trafficking patterns. Yolk in *C. elegans* consists of both lipids and proteins (*69*). Because low lipid levels in *sbp-1(RNAi)* animals makes results with VIT-2::GFP difficult to interpret, we used the *myo-3*::ssGFP reporter model (*61*), which expresses and secretes GFP in body wall muscle cells.

We conducted a small scale screen of 19 other STRESS RESPONSE: Pathogen: unassigned genes on secretion in *myo-3*::ssGFP and the sensitized *sbp-1(ep79)*; *myo-3::GFP* backgrounds, imaging body wall muscle cells in triplicate and manually scoring aggregation in blinded samples (**Fig6A**). *hpo-6* and two other candidate genes, *irg-2* and ZK6.11/*irg-8* were validated by confocal imaging, blinded visual scoring and quantitation of puncta size and number with Cell Profiler (*70*). We found that as in *arf-1* RNAi, reduction in *hpo-6, irg-2* and *irg-8*/ZK6.11 increased puncta size (**Fig6B-D; FigS6; D-E**), with related decrease in aggregate number in both wild type and the *sbp-1(ep79)* background, reflecting aggregation and disruption of normal secretory processes.

**Figure 6:**
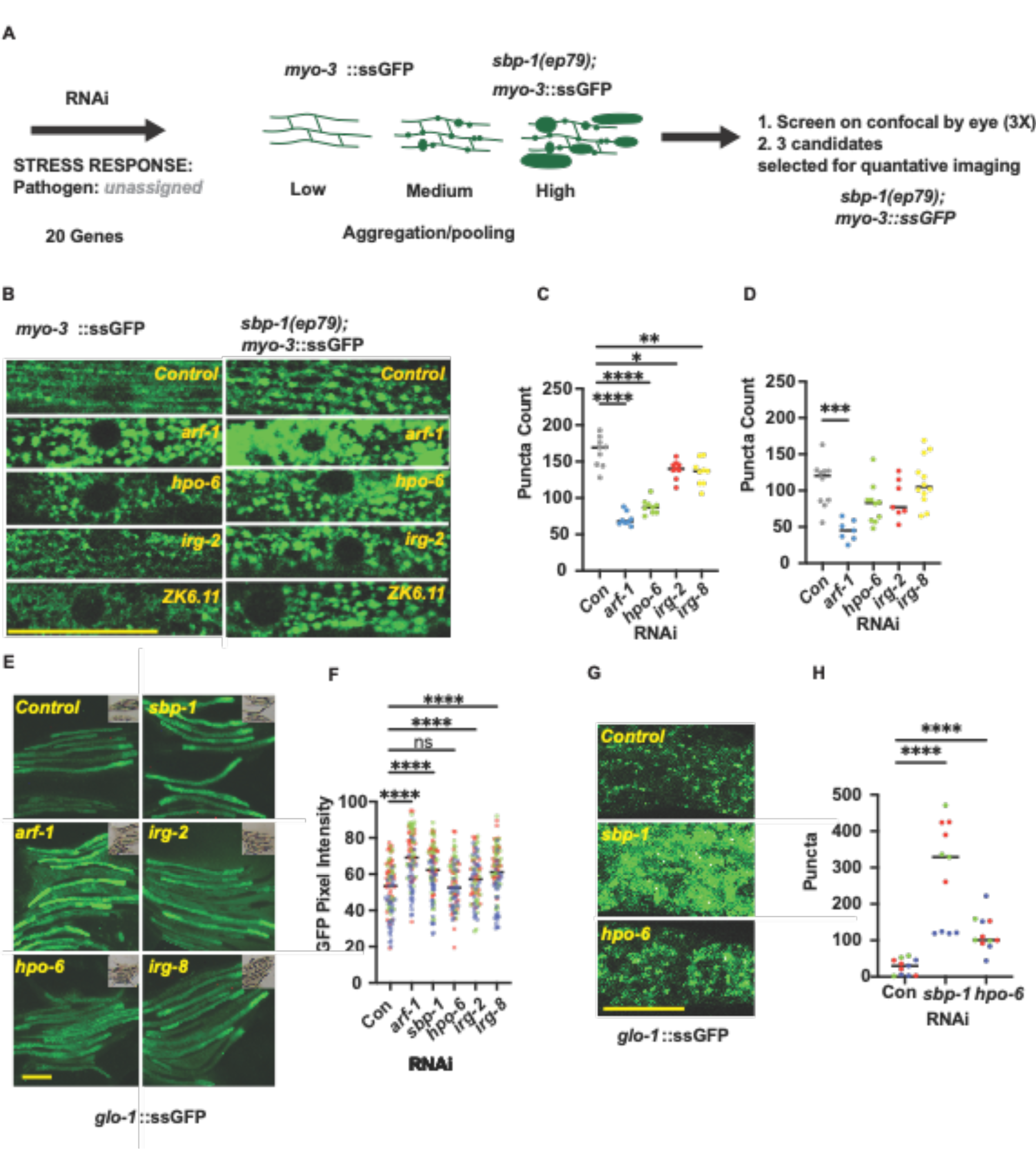
ILG loss can enhance aggregation of secreted proteins. **A.** Schematic diagram of small-scale screen of STRESS RESPONSE: Pathogen: *other* genes for aggregation of a secreted reporter in body wall muscle (*myo-3*::ssGFP). Confocal projections of ssGFP in body wall muscle in control or the *sbp-1(ep79)* background (**B**). Quantification of three independent replicates by Cell Profiler determining number of puncta (**C, D**) and puncta area (**Figure S6D, E**). **E**. Epifluorescence images of ssGFP expressed from an intestinal reporter with DIC images as insets. Quantitation of pixel intensity with ImageJ is in **F**. Confocal projections showing ssGFP aggregation in **G**, with puncta quantitation by Cell Profiler in **H**. Significance is determined by the Mann-Whitney test.

Many of the STRESS RESPONSE: Pathogen genes are expressed in the intestine; therefore, we also examined a ssGFP expressed under the *glo-1* promoter (*71*) (**Fig6E**). Using epifluorescence imaging, we found that intestinal ssGFP accumulated after RNAi of *sbp-1, arf-1, hpo-6, irg-2* and *irg-8* (**Fig6F, G**). Confocal imaging shows that ssGFP puncta aggregate in the intestine of *sbp-1(RNAi)*animals, as in muscle tissue and that this also occurs with *hpo-6* RNAi (**Fig6G, H**). Thus, in a variety of cell types, we find that interference with *sbp-1*, or candidate STRESS RESPONSE: Pathogen genes, causes aggregation and pooling of ssGFP in the absence of pathogen stimulation, suggesting a role in secretory processes.

### Knockdown of STRESS RESPONSE: Pathogen genes affects an endogenous PA14 stimulated peptide

The secretory pathway carries out many basal functions and must increase capacity during stresses which require increases in protein production, such as pathogen challenge. The ER stress pathway plays a key role during this increase (*72, 73*) and our results suggest that alterations in the Golgi may also be sensed and result in upregulation of ILGs. Our studies ssGFP reporters allowed isolation of effects of lipid imbalances on basal secretory processes from the complex response to pathogens. However, ILGs are defined by high level expression upon pathogen exposure and could also affect secretion of antimicrobial peptides (AMPs) or other defensive molecules. *C. elegans* contains a large number of proteins with domains common to AMPs such as ShKT, Sushi and CUB domains (*74*). In order to test effects of the ILGs on an endogenous secreted protein in the pathogen response, we selected C32H11.9/*cld-1* (cub-like domain 1). In our previous RNA seq experiments, it was the highest upregulated gene after PA14 exposures (**Fig7A**) and was also strongly induced during other early timepoints (*22*), but not induced by lipid imbalances in *sbp-1(RNAi)* animals (**Fig7B**). It contains a signal sequence, suggesting that is secreted and a CUB domain (**Fig7C**). We obtained an allele with GFP fused to the C terminus of *cld-1*/C32H11.6 using CRISPR (PHX2081(C32H11.6::GFP(*sybIS7080*)), which accumulated in intestine after PA14 exposure (**Fig7D**), consistent with disruption with secretion. Notably, RNAi of *sbp-1* disrupted CLD- 1::GFP production, causing increases total levels, aggregation and accumulation in the pseudocoelom, consistent with defects in secretory processes resulting from lipid imbalances (**Fig7D**). Knockdown of with *arf-1* also produced similar effects, strengthening these links. Interference with *hpo-6*, a candidate ILG, also caused accumulation and aggregation of CLD-1::GFP during PA14 exposure, suggesting that some ILG genes act to support secretory function during stress, which may be initiated by lipid imbalances or pathogen exposure.

**Figure 7:**
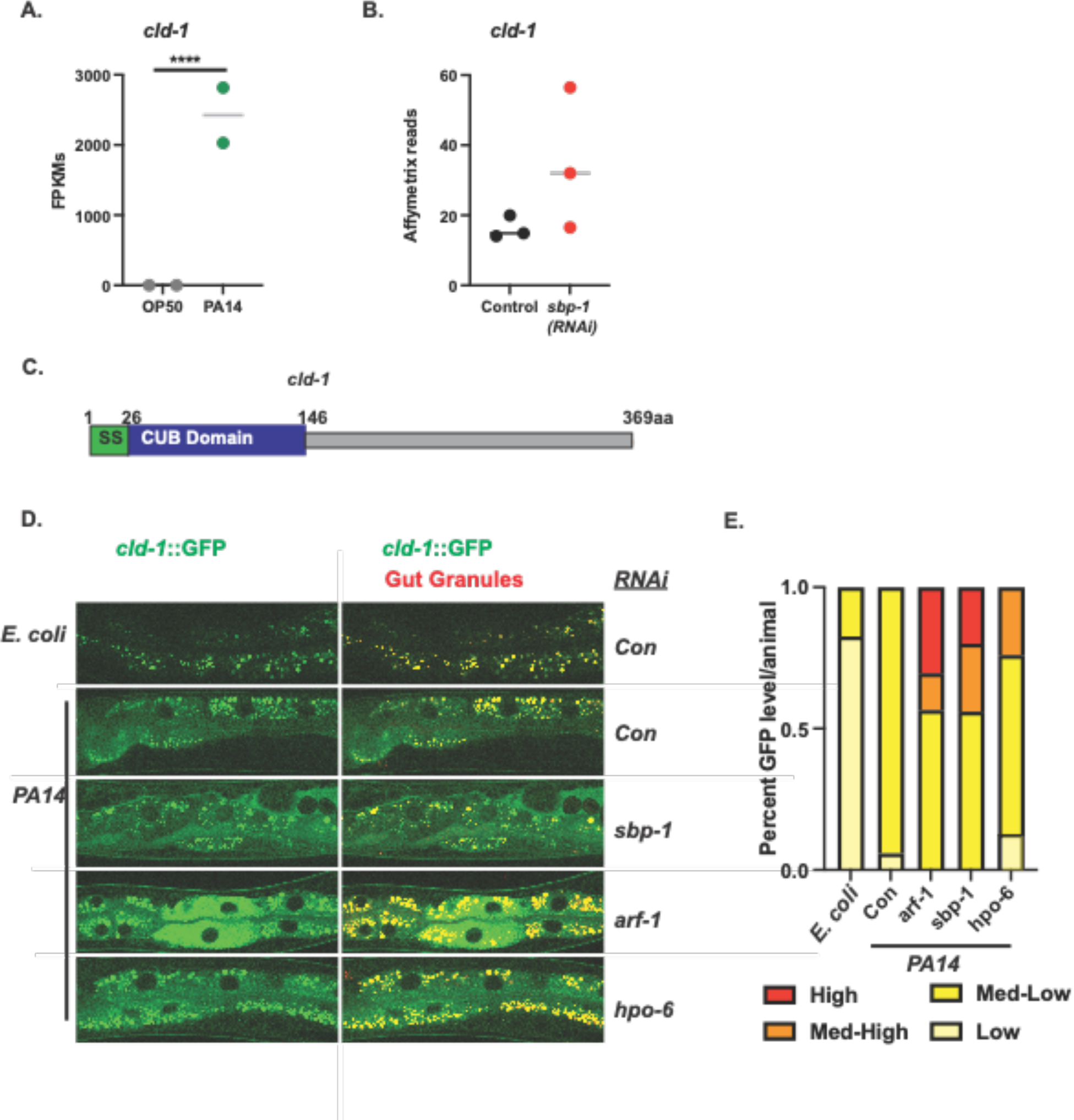
ILG loss can enhance intestinal accumulation of a Pseudomonas-induced CUB domain peptide. Expression levels of C32H9.11 in response to Pseudomonas aeruginosa (PA14) from Ding, et al. 2018 (**A**) or after *sbp-1* RNAi from Ding, et al. 2015 (**B**). **C.** Domain structure of C32H11.9 showing signal sequence and CUB domain. **D.** Spinning disc confocal micrographs showing intestinal accumulation of *cld-1*/C32H11.9::GFP in response to PA14 after *sbp-1*, *arf-1* or ILG RNAi taken at 488mn. Gut granules are shown with an overlay of a micrograph of the same section from taken at 564nm. Quantitation is in **E**.

## Discussion

Pathogen elicited gene expression programs are part of a specialized stress response where specific sets of antimicrobial peptides are produced, then secreted in response to pathogens. This type of innate immune response initiates as pathogens are recognized by interaction with cellular proteins or cellular processes are disturbed (*75*). In *C. elegans*, evidence for microbe/pathogen associated molecular patterns (MAMPS/PAMPS) has been less clear, however, surveillance of cellular processes such as translation (*11*), initiating the unfolded protein response or impacting lipid bilayer function activates pathogen responsive signaling and transcription of ILGs (*9, 15, 16, 34*). While some studies have suggested changing levels of specific lipids can prime pathogen responses (*34*), lipid perturbations activating pathogen-responsive gene expression do not necessarily confer pathogen resistance (*9*), and ILG expression levels may be much lower than what occurs in a pathogen response. Links between membrane lipid levels and immune gene activation also occur in other organisms. In *Drosophila*, synthesis of PC is critical to immune function, balancing stored lipids with need to maintain membranes for secretory processes (*76*). In mammals, interference with a master transcriptional regulator of lipid biosynthetic enzymes, *sbp-1/SREBP-1* activates a gene expression program shared with pathogen stress in both *C. elegans* and mammals (*14*). This raises the larger the question of why ILG expression increases when membrane lipids are unbalanced in the absence of pathogens.

Membrane lipids form the organelles of the secretory system, which can sense changes in protein homeostasis or lipids levels and elicit gene expression programs when stressed. Multiple well-studied pathways are linked to changes in proteostasis or lipid levels within the ER (*57*). These processes are orchestrated by ER-intrinsic proteins that sense changes in protein folding or alterations in the lipid bilayer (*6, 8, 15, 77*). These ER-linked events are also important during pathogen exposure in *C. elegans*; changes in ribosomal function at the ER have a well-defined impact on pathogen responses (*11, 78*) and pathogen responses depend on ER stress pathways to protect this organelle from the demands of the immune response (*73, 79*). Interestingly, the Lee lab also found that ILGs are upregulated when trafficking is inhibited through blocking glycosylation in the ER (*80*). In addition, pathogen-responses require an intact ER stress pathway to allow adjustment to the increased trafficking load (*58, 73*), as antimicrobial peptides are produced in mass in the ER, sent in lipid vesicles to the Golgi, where many are glycosylated and then packaged into lipid vesicles for secretion (*81*). For example, the *C. elegans* genome contains more than 300 genes that are produced in response to specific pathogens and which share similarity to known antimicrobial effectors (*82*). However, many of the genes in the *C. elegans* innate immune response do not have obvious similarity to antimicrobial effectors and are defined solely as being responsive pathogens (*27, 83*). It is this class that are largely shared between pathogen-induced and membrane stress-induced stress programs. Several lines of evidence prompted us to hypothesize that some of these genes might be part of a response linked to stress in membranes of the secretory pathway rather than a direct response to an extrinsic pathogen. First, the involvement of secretory organs was implicated by alterations in PC ratios in the ER/Golgi in both *sams-1* and *sbp-1* RNAi lipidomes. Second, our targeted RNAi screen for regulators of p*sysm-1::GFP* identified the *arf-1* GTPase and coatomer proteins, which are critical Golgi/ER transport. Importantly, knockdown of *arf-1* was sufficient to induce ILG upregulation. Strengthening this link between disruption of lipid levels and ARF-1 function, we also found *sbp-1* RNAi altered ARF-1::mCherry localization and Golgi morphology.

Other organelles in the secretory pathway, such as the Golgi, have their own stress sensors and response pathways (*84*), some of which act by impacting the ARF1 GTPase (*85*). ARF-1 is a critical regulator of trafficking, coordinating retrograde traffic from the Golgi to the ER and regulating secretory function in the trans-Golgi (*86*). Cycling of ARF-1 GTPases depends on membrane localization of the ARF GTPase, GEF (Guanine exchange factor), and GAP (GTPase activating protein) (*86*). We previously found that knockdown of PC synthesis enzymes blocked ARF1 cycling and limited membrane association of the ARF GEF GBF1 in cultured human cells (*25*). This suggests low PC levels affect ARF1 activity by limiting the ability of the GTPase, GEF, and GAP to associate at the Golgi membrane and initiate GDP-GTP cycling. However, PC could be linked to ARF1 through other mechanisms. ARF1 and phospholipase D (PLD) function have been linked by multiple labs (*87*). PLD cleaves PC molecules to produce choline and phosphatidic acid, which in turn stimulates vesicle formation (*88*). This regulatory loop requires PC, which is low in ER/Golgi membranes in *sams-1* and *sbp-1* RNAi animals. DAG kinases, which could be stimulated by changes in DAGs in the *sbp-1* or *sams-1(RNAi)* animals, can also affect ARF activity and trafficking (*89*). Finally, Protein Kinase D has an important role in maintaining Golgi structure, and its ortholog has been linked to immunity in *C. elegans* (*90, 91*). However, DAG kinase or PKD (Protein Kinase D) loss failed to activate the p*sysm-1::GFP* reporter in our RNAi screen, suggesting they could be important in other regulatory contexts. While the lipidomic changes in *sbp-1* and *sams-1* animals are broad and individual lipids could act through multiple independent mechanisms, our results show that decreasing PC is sufficient to limit ARF-1 function (*25*) and activate ILG expression in the absence of pathogen.

Stress-responsive genes, such as ILGs, may contain several classes of factors with distinct functions. Many of the ILGs, for example, those in the STRESS RESPONSE: Pathogen: unassigned WormCat categories (*27, 28*), lack domains associated with AMP function. While some could have antimicrobial properties with distinct motifs, the activation of these genes through intrinsic stress in the Golgi prompted us to ask if others acted to support secretory function. Our data showing that reduction of *hpo-6, irg-2* and *irg-8* increased aggregation and pooling of a secreted reporter in control and *sbp-1(ep79)* backgrounds suggests they could protect or support secretory function occurring after broad disruption in membrane lipids or after pathogen exposure occurs because both processes stress the trafficking system. Thus, some ILGs may function generally a “multi-membrane” stress response encompassing the roles of both the ER and the Golgi in trafficking. Taken together, these results illustrate mechanisms linking lipid metabolism to genes activated in the immune response through effects on the secretory system.

## Materials and Methods

### *C. elegans* strains, RNAi constructs, and screening

N2 (wild type), p*sysm-1::GFP (*AU78*), myo-3*::GFP (GS1912), MANS::GFP (RT1315) and OP50-1 were obtained from the *Caenorhabditis* Genetics Center. Exglo-1::ssGFP (GH639) was a gift from Dr. Greg Hermann (Lewis and Clark University). *myo-3::ssGFP*;*sbp- 1(ep79)*(WAL510) was constructed in this study. CRISPR-tagged ARF-1, p*arf-1::ARF- 1::mCherry (knu418),* was obtained from the In Vivo Biosystems (COP1415) and then outcrossed 3 times (WAL10). Imaging of MANS::GFP was done from a strain also harboring a CRISPR tagged warf-1::RFP (ker4) (WAL12). (*cld-1*/C32H11.6::GFP was constructed by SUNY Biotech using CRISPR (PHX2081(*sybIS7080*)). Normal growth media (NGM) was used unless otherwise noted. For RNAi, gravid adults were bleached onto NGM plates supplemented with ampicillin, tetracycline, and 6mM IPTG and 10X concentrated bacterial cultures. *C. elegans* were allowed to develop until the young adult stage before harvesting. Because of larval arrest, *copa-1* RNAi bacteria were diluted 1:10 in control RNAi bacteria before plating. For the RNAi screen of *psysm-1*::GFP, L1 larvae were plated into 96 well plates spotted with RNAi bacteria, and L4/young adults were scored from -3 to +3 with 0 as no change in 4 independent replicates on a Zeiss GFP dissecting scope with an Axiocam camera. Candidates that were positive in ¾ replicates were tested an additional four times, imaging 3 sets of 10 animals per RNAi. Images quantitation is described below. For RNAi screen of *myo-3*::ssGFP and *myo- 3*::ssGFP; *sbp-1(ep79)*, 4 independent replicates were subjected to RNAi, placed in channeled agarose pads and confocal images were collected on a Leica SP6.

### Cell culture and siRNA transfection

HepG2 cells (ATCC, HB-8065) were grown in Minimum Essential Medium (Invitrogen) plus 10% FBS (Invitrogen), Glutamine (Invitrogen), and Sodium Pyruvate (Invitrogen). siRNA oligonucleotides transfections were done for 48 hours using Lipofectamine RNAiMAX Transfection Reagent (Invitrogen, 13778100), and cells were held for 16 hours in 1% Lipoprotein Deficient Serum (LDS) (Biomedical Technologies, BT907) and 25 μg/ml ALLN (Calbiochem) for 30 min prior to harvesting.

### Immunoblotting

**HepG2 cells:** Syringe passage was used to lyse cells in High Salt RIPA (50 mM Tris, pH 7.4; 400 mM NaCl; 0.1% SDS; 0.5% NaDeoxycholate; 1% NP-40; 1 mM DTT, 2.5 μg/mL ALLN, Complete protease inhibitors (Roche). Invitrogen NuPage gels (4- 12%) were used for protein separation before transfer to nitrocellulose. Blots were probed with antibodies to phosphorylated p38 MAP Kinase, total p38, and STAT1 were used as a control. Immune complexes were visualized with Luminol Reagent (Millipore). Densitometry was performed by scanning the film, then analysis of pixel intensity with ImageJ software. Graphs show the average of at least three independent experiments with control values normalized to one.

***C. elegans:*** Young adult animals were lysed by sonication in High Salt RIPA, and immunoblotting was performed as above.

### Microscopy and Image quantitation

ARF-1::mCherry, MANS-1::GFP and ssGFP images were acquired on a Leica SPE confocal, and projections of confocal slices were produced. All images were taken at identical gain settings within experimental sets, and Adobe Photoshop was used for corrections to levels across experimental sets. Image quantitation of GFP fluorescence was done using FIJI/ImageJ2. Briefly, each photo was processed individually, beginning by splitting the image into three color channels (Red/Green/Blue) and choosing the green channel for analysis. This image was then converted to an 8-bit grey image and processed using the function “Enhance Local Contrast (CLAHE)” (Blocksize: 250, Histogram bins: 256, Max Slope: 5.00, Mask: none, Fast (less accurate): no (unchecked)). Using the circle selection tool, four sections (w: 55 h: 25) of the brightest areas across each worm’s intestine were selected from this processed image and measured for intensity (integrated intensity). One additional section was taken from beside each worm to account for background fluorescence. The background reading was subtracted from each individual intestinal reading, then the four corrected readings were added to get a sum of fluorescent intensity. Imaging of C32H11.9/*cld-1*::GFP was performed on a W1 Yokogawa spinning disk confocal microscope equipped with a Photometrics Prime BSI Express sCMOS camera. Quantitation was performed with Nikon Imaging System software. Statistical tests performed include Normality and Lognormality tests (Anderson-Darling, D’Agostino & Pearson, Shapiro-Wilk, Kolmogorov-Smirnov) and Mann-Whitney tests.

### Lipidomics

*C. elegans* total and microsomal lipidomics, including fractionation protocols, were performed at the Whitehead Metabolomics core as in Smulan et al. 2016. Statistical analysis was performed in Graphpad prism.

### Gene expression analysis

Lysis of young adult *C. elegans* was performed in 0.5% SDS, 5% β-ME, 10 mM EDTA, 10 mM Tris-HCl ph7.4, 0.5 mg/ml Proteinase K, before purification of RNA by TRI-Reagent (Sigma). cDNA was produced with Transcriptor First-strand cDNA kits (Roche), and RT-PCR was performed using Kappa SYBR Green 2X Mastermix.

RNA for sequencing was purified using RNAeasy columns (Qiagen). RNA sequencing (including library construction was performed by BGI (Hong Kong). Reads were analyzed through the Dolphin analysis platform (https://dolphin.umassmed.edu/), using DeSeq to identify differentially expressed genes (*92*). Gene set enrichment was performed using WormCat (www.wormcat.com) (*27, 28*).

## Acknowledgments

We would like acknowledge Drs. Seung-Jae Lee, Sujeong Kwon (Korea Advanced Institute of Science and Technology) for providing the *lpin-1* RNA seq data. In addition, Dr. Greg Hermann provided the *glo-1*::ssGFP strain (GH639). We would also like to thank Drs. Seung-Jae Lee, Emily Troemel (USD), Cole Haynes (UMASS Chan School of Medicine), Alexandra Byrne (UMASS Chan School of Medicine) and members of the Walker lab for comments on the manuscript in addition to Dr. Marian Walhout (UMASS Chan School of Medicine) for helpful discussion. We would like to acknowledge Wei Ding for technical support and Caroline Lewis (MIT) for metabolomics screening at the Whitehead Metabolomics core. We also thank Life Science Editors and Dr. Sabbi Lall for manuscript editing. Funding was from the NIA at the NIH: R01AG053355, R56AG068670 and R01AG068670. Some strains were provided by the CGC, which is funded by the NIH Office of Research Infrastructure Programs (P40 OD010440).

## Author Contributions

Conceptualization; AKW, MJF, CMW, LJS

Verification: AKW, MJF, CMW, LJS

Formal Analysis: AKW, MJF, CMW, LJS, DML

Investigation: AKW, MJF, CMW, LJS, DL, MF

Resources: AKW

Data Curation: AKW

Writing: AKW

Writing/Editing: AKW, MJF, CMW, DSL, LJS

Visualization: AKW

Supervision: AKW

Funding: AKW

## Data and materials availability

All data are available in the main text or the supplementary materials will be shared. GEO submission of RNA seq data is in progress.

