## Supplemental figures and legends for "Stimulation of immunity-linked genes by membrane disruption is linked to Golgi function and the ARF-1 GTPase"

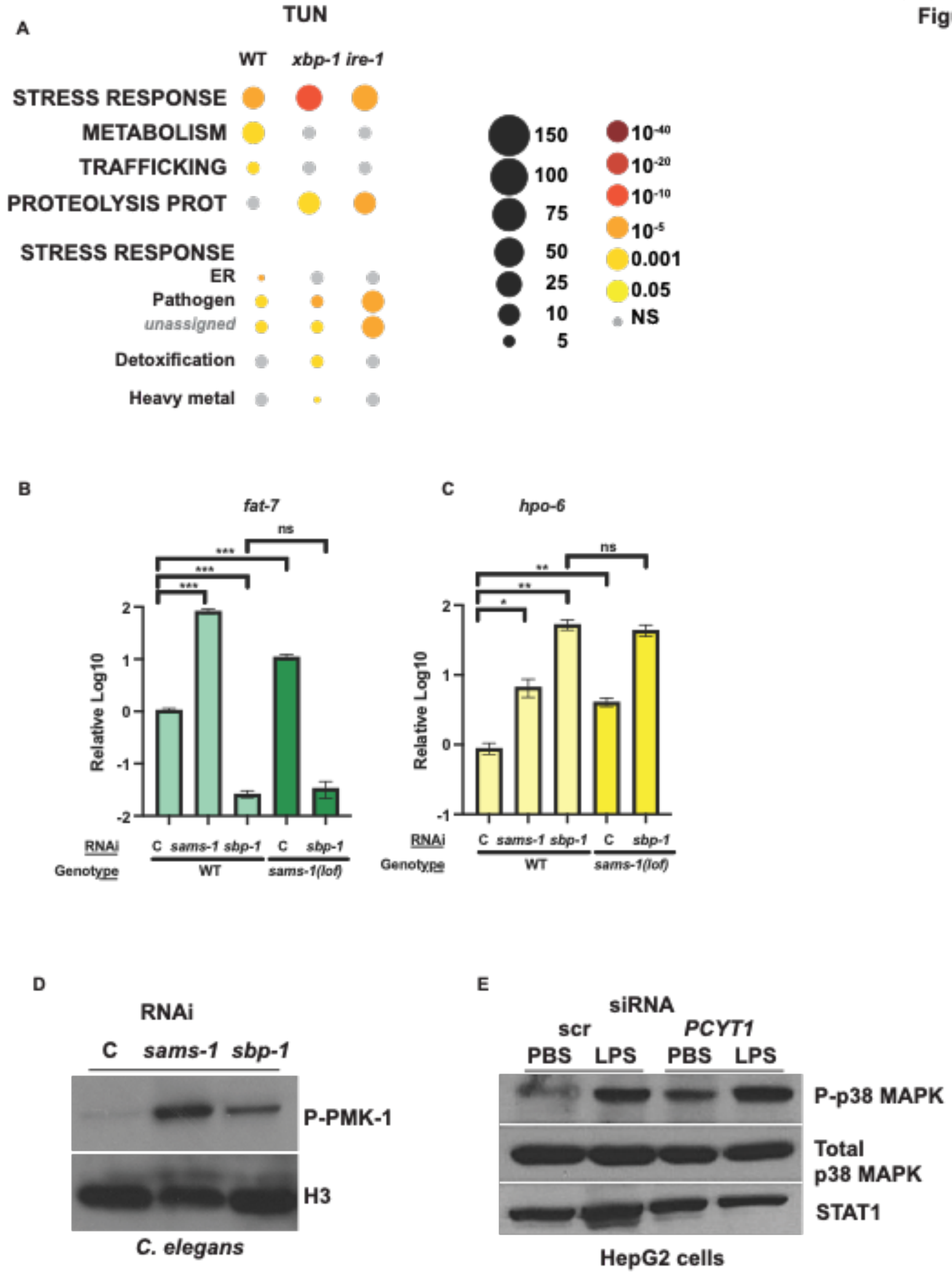


**Fig. S1.**

**Figure S1: Activation of Stress response pathways when PC synthesis is disrupted. A.**

WormCat enrichment analysis of Shen, et al. comparing control and ER stress RNAi treatments after tunicamycin treatment. **B.** qRT-PCR shows that upregulation of an immunity-linked gene, *hpo-6*, in *sams-1(lof)* animals is not dependent on *sbp-1* (**C**). **D.** The p38 MAP kinase MAPK14 is phosphorylated after siRNA of *PCYT1* in human hepatoma cells. **E.** Constitutive phosphorylation of the *C. elegans* MAPK14 ortholog, PMK-1, occurs after both *sams-1* and *sbp-1(RNAi)*.

A

| | Class total | *sams-1* | *sbp-1* |
|---|---|---|---|
| PE | 376 | 116 | 134 |
| PC | 295 | 125 | 122 |
| TAG | 154 | 78 | 67 |
| LPE | 106 | 6 | 9 |
| PS | 85 | 9 | 22 |
| LPC | 72 | 9 | 8 |
| DG | 56 | 21 | 14 |
| Cer | 53 | 8 | 16 |
| CL | 50 | 4 | 13 |
| PG | 47 | 3 | 4 |
| PI | 44 | 0 | 5 |
| PA | 42 | 1 | 0 |

Species Distribution
>0.05
>0.0001

B

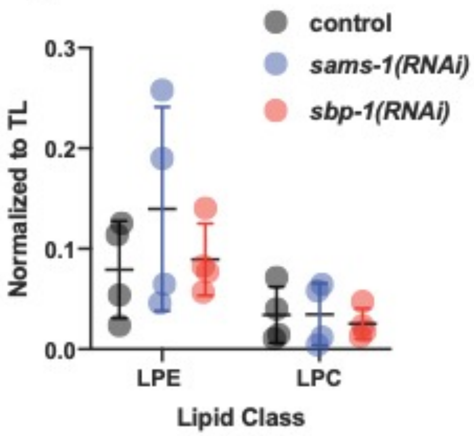


C

| | Class Total | *sbp-1* |
|---|---|---|
| PE | 355 | 128 |
| PC | 312 | 156 |
| TAG | 83 | 44 |
| LPE | 66 | 7 |
| PS | 102 | 39 |
| LPC | 54 | 6 |
| DG | 77 | 47 |
| Cer | 38 | 11 |
| CL | 52 | 28 |
| PG | 101 | 3 |
| PI | 57 | 19 |
| PA | 49 | 8 |
| SM | 33 | 11 |

Species Distribution
>0.05
>0.0001

D

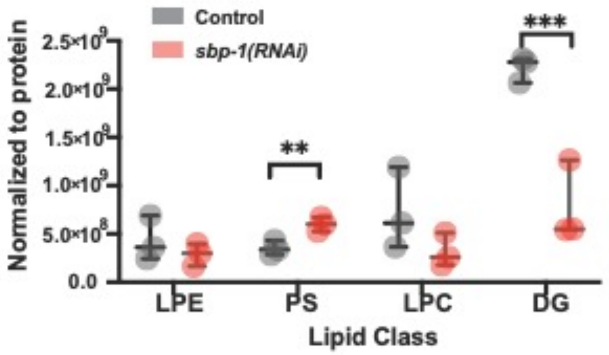


**Fig. S2.**

**Figure S2: Broad lipidomic changes in total and microsomal lipid compartments after *sams-1* or *sbp-1* RNAi**. Tables showing numbers lipid species within each class in the total lipidome (**A**) or ER/Golgi lipidome in *sbp-1(RNAi)* animals (**B**). Green boxes denote classes in which the distribution of the species within each RNAi significantly differed from controls using the Wilcoxen test for non-parametric data. Levels of lipid classes in total *sams-1* or *sbp-1* total lipidomics (**D**) or *sbp-1* ER/Golgi lipidomics (**D**).

**Fig. S3.**

Fanelli, et al.
Figure S3

| Gene ID | Gene name | *fat-7*::GFP | WormCat Category |
|---|---|---|---|
| B0336.2 | *arf-1* | up | Trafficking: ER/Golgi: ARF |
| Y6B3A.1 | *agef-1* | | Trafficking: ER/Golgi: ARF |
| F59E10.3 | *copz-1* | | Trafficking: ER/Golgi: COP I |
| Y71F9AL.17 | *copa-1* | * | Trafficking: ER/Golgi: COP I |
| Y25C1A.5 | *copb-1* | * | Trafficking: ER/Golgi: COP I |
| T14G10.5 | *copg-1* | * | Trafficking: ER/Golgi: COP I |
| ZK180.4 | *sar-1* | * | Trafficking: ER/Golgi: COP II |
| C23H3.4 | *sptl-1* | | Metabolism: lipid: sphingolipid |
| T10H9.5 | *pmp-5* | | Peroxisome: transporter |
| Y37E11AR.4 | *nape-1* | * | Signaling: lipid: phospholipase: other |
| B0285.10 | *ckb-3* | up | Metabolism: lipid: phospholipid |
| C39D10.3 | *pcyt-2.2* | | Metabolism: lipid: phospholipid |
| Y22D7AL.8 | *sms-3* | up | Metabolism: lipid: sphingolipid |
| F53H8.4 | *sms-2* | | Metabolism: lipid: sphingolipid |
| F32D8.4 | | | Trafficking: ER/Golgi: Golgi other |
| F55A11.2 | *syx-5* | | Trafficking: vesicle targeting/fusion: SNARE |
| W02B12.1 | | | Signaling: lipid: phospholipase B |
| T23H2.5 | *rab-10* | | Trafficking: vesicle targeting/fusion: RAB |
| ZC506.3 | *pssy-1* | up | Metabolism: lipid: sphingolipid |
| F38E11.5 | *copb-2* | | Trafficking: ER/Golgi: COP I |
| W06D12.3 | *fat-5* | nd | Metabolism: lipid: fatty acid |
| F10D2.9 | *fat-7* | nd | Metabolism: lipid: fatty acid |
| VZK822L.1 | *fat-6* | nd | Metabolism: lipid: fatty acid |

**Screen hits**

**Selected library genes included in rescreen**

**Other lipid metabolism genes included in rescreen**

**Figure S3: Candidates from lipid sub-library RNAi screen.** List showing candidates from quadruplicate screening of the RNAi sub-library along with other genes selected as controls for the rescreen.

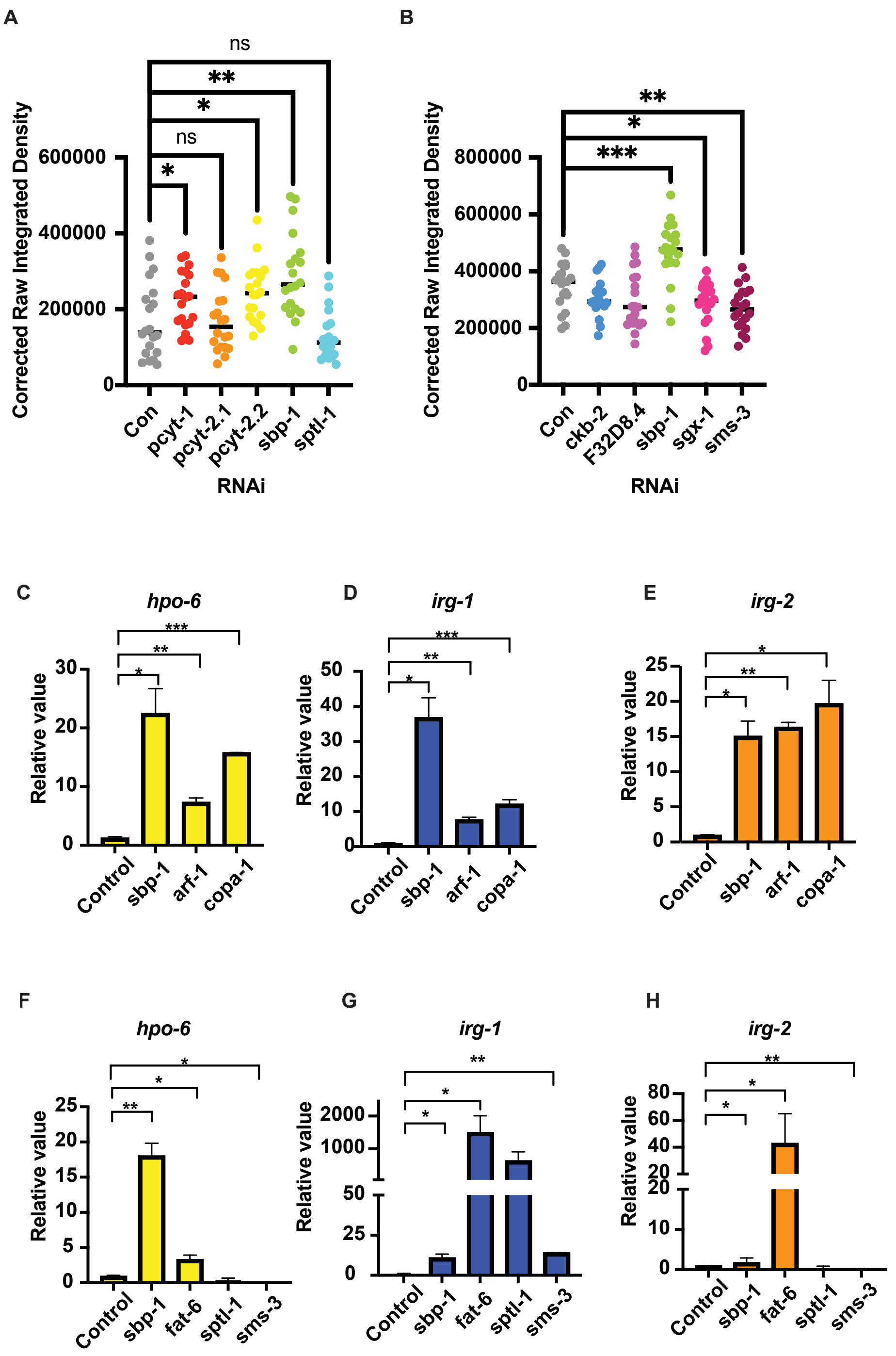


**Figure S4: Regulation of immunity-linked genes in conditions of membrane disruption.** Quantitation GFP intensity after knockdown of Golgi trafficking regulators **(A)** or lipid synthesis genes **(B)** as calculated by ImageJ. Because the data did not follow normal distribution patterns, the Mann-Whitney test was used to calculate significance. qRT-PCR from *psysm-1*::GFP (**C-E**)

or p*sysm-1*::GFP; *sbp-1(ep79)* (**F-H**) shows effects of disrupting *arf-1* or membrane lipids on ILGs. Significance is calculated by Students T Test. Error bars show standard deviation.

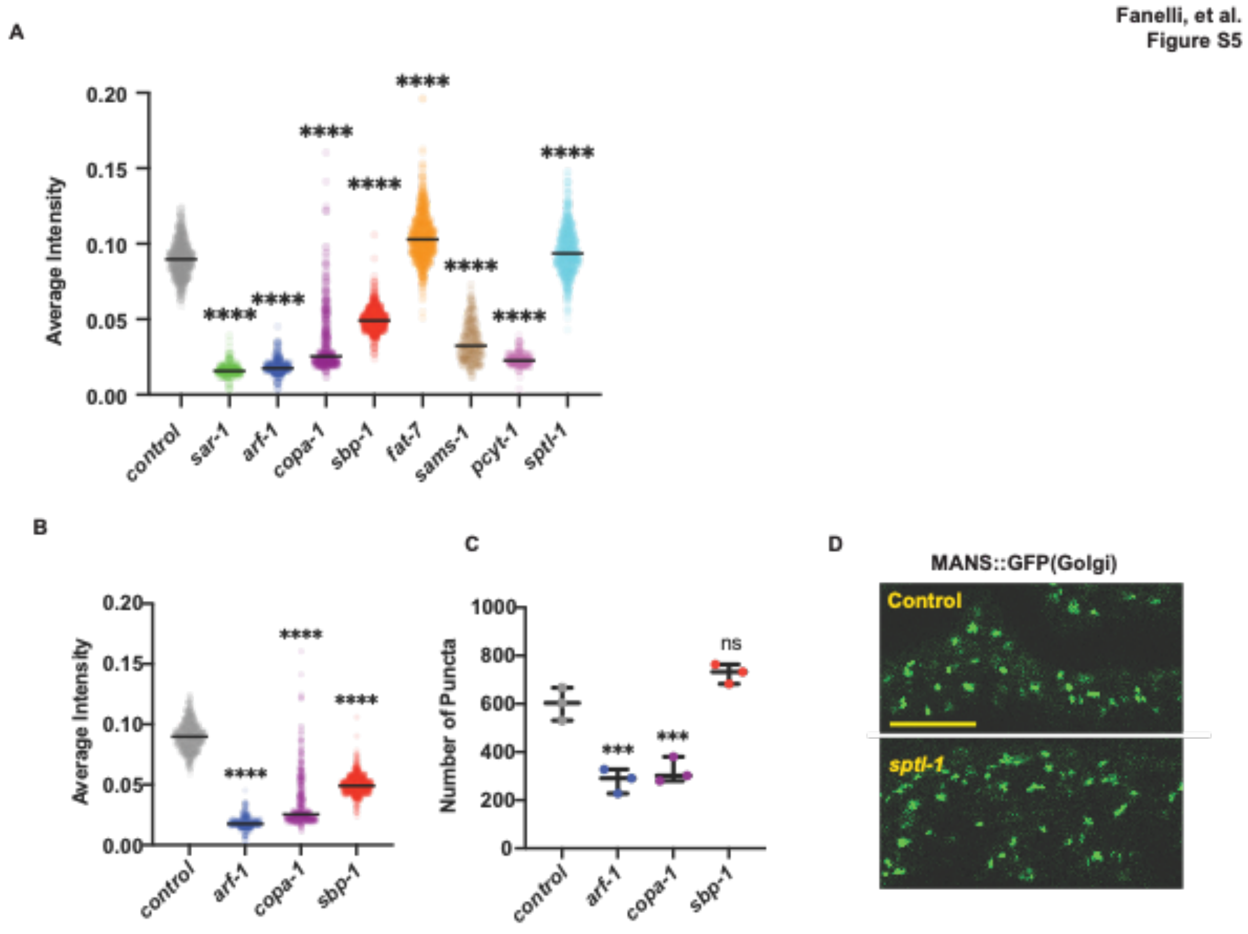


**Figure S5: Shared upregulation of immunity linked genes in *sams-1, sbp-1* and *arf-1* transcriptomic studies.** A. Quantitation of epifluorescence images from Figure 3 D-J calculated by ImageJ. Because data did not follow normal distribution patterns, Mann-Whitney test was used to calculate significance. Quantitation of puncta intensity (**B**) and number (**C**) from ARF-1::mCherry images in Figure 4G measured by Cell Profiler. Significance calculated by Mann Whitney. **D.** Confocal micrographs showing minimal changes after *sptl-1* RNAi in of MANS::GFP. Bar show 10 microns.

Fanelli, et al.
Figure S6

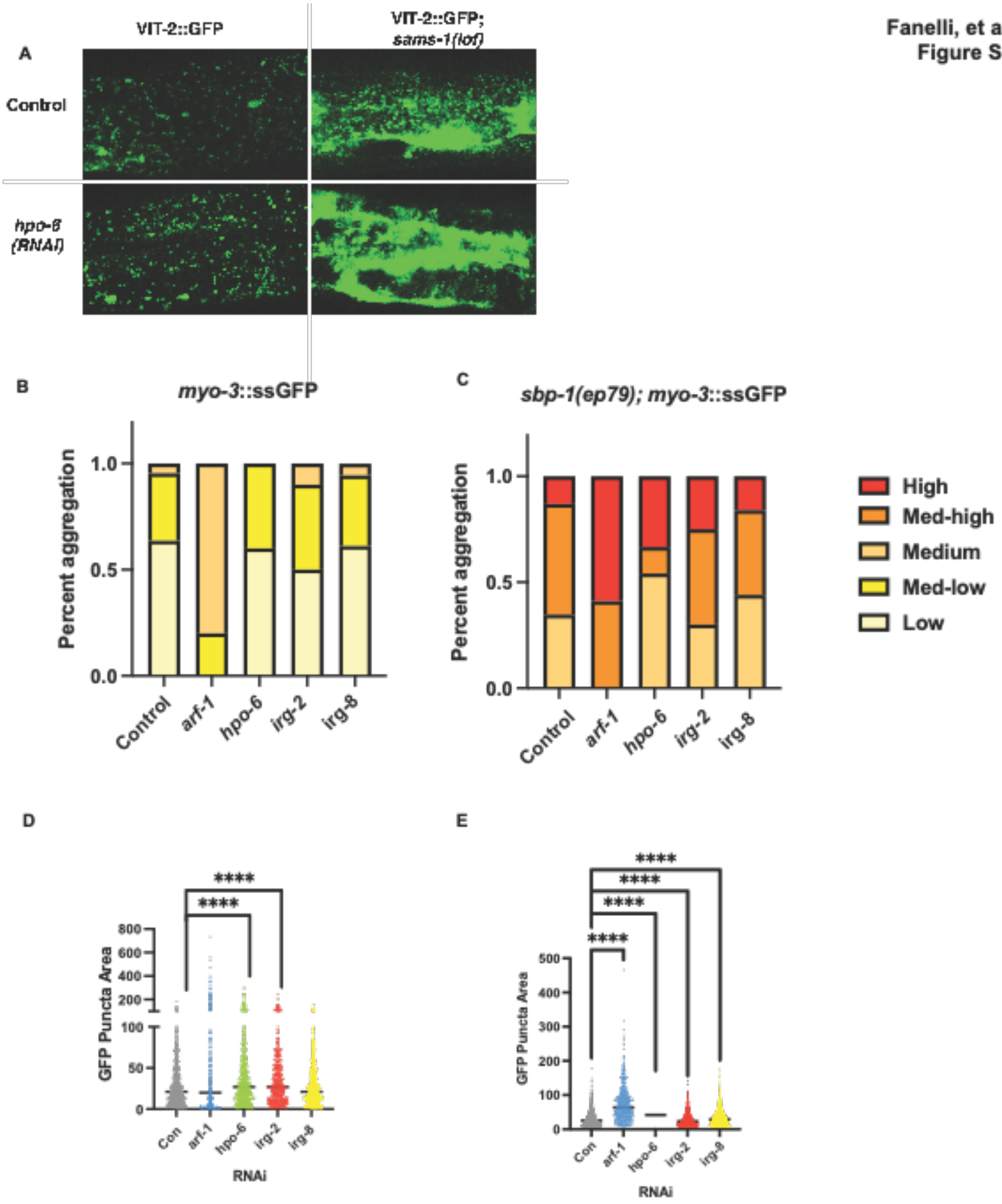


**Figure S6: Reduction in *hpo-6* exacerbates trafficking defects**. A. Confocal projections of *C. elegans* intestinal cells expressing *vit-2*::GFP (RT130) in wild type or *sams-1(lof)* backgrounds. B, C. Visual scoring of confocal images from Figure 6B. Scoring was done blind on three independent experiments. Quantitation of GFP puncta area from Figure 6B (D, E).

**Data S1: Transcriptomics and gene set enrichment from *lpin-1(RNAi)* animals.** 2 fold upregulated genes in *lpin-1(RNAi)* animals. Gene set enrichment was performed using WormCat (*27*).

**Data S2**: **Total lipidomics of *sams-1* and *sbp-1(RNAi)* animals and microsomal lipidomics of *sbp-1(RNAi)* adults.** Datasets from two lipidomic studies. The first is LCMS data normalized to total lipids compared between Control, *sbp-1,* and *sams-1*(*RNAi)* adults. Tabs are labeled sams_sbp_Lipid class. The second dataset contains LCMS from Control and *sbp-1(RNAi)* microsome preps from adult animals. Significance of fold changes in individual lipids determined by students T-test.

**Data S3: Screen of lipid sub-library with an immune reporter.** Tab 1 contains averaged results from quadruplicate screening of p*sysm-1*::GFP. The secondary screen contains averages from candidates selected from the first screen, plus *fat-5, fat-6,* and *fat-7* as additional test RNAis. The secondary screen was conducted in quadruplicate. ND is not done.

**Data S4**: **Transcriptomic responses to trafficking disruption.** RNAseq from control vs. *arf-1/arf-1(RNAi)* animals (arf_1_all, arf_2Xup) and gene set enrichment analysis produced in WormCat for Category 1, Category2 and Category3). WormCat analysis of *vit-2*::GFP (*61*) and *glp-1* (*60*) datasets.
